# The art of seeing the elephant in the room: 2D embeddings of single-cell data do make sense

**DOI:** 10.1101/2024.03.26.586728

**Authors:** Jan Lause, Philipp Berens, Dmitry Kobak

## Abstract

A recent paper in *PLOS Computational Biology* (Chari and Pachter, 2023) claimed that *t*-SNE and UMAP embeddings of single-cell datasets fail to capture true biological structure. The authors argued that such embeddings are as arbitrary and as misleading as forcing the data into an elephant shape. Here we show that this conclusion was based on inadequate and limited metrics of embedding quality. More appropriate metrics quantifying neighborhood and class preservation reveal the elephant in the room: while *t*-SNE and UMAP embeddings of single-cell data do not preserve high-dimensional distances, they can nevertheless provide biologically relevant information.

---

In single-cell genomics, researchers often visualize data with 2D embedding methods such as *t*-SNE (Van der Maaten and Hinton, 2008; Kobak and Berens, 2019) and UMAP (McInnes et al., 2018; Becht et al., 2019). Chari and Pachter (2023) criticize this practice: They claim that the resulting 2D embeddings fail to faithfully represent the original high-dimensional space, and that instead of meaningful structure these embeddings exhibit “arbitrary” and “specious” shapes. While we agree that 2D embeddings necessarily distort high-dimensional distances between data points (Nonato and Aupetit, 2018; Wang et al., 2023b), we believe that UMAP and *t*-SNE embeddings can nevertheless provide useful information. Here, we demonstrate that UMAP and *t*-SNE preserve cell neighborhoods and cell types, and that the conclusions of Chari and Pachter (2023) are based on inadequate metrics of embedding quality.

To illustrate their point that *t*-SNE and UMAP embeddings are arbitrary, Chari and Pachter (2023) designed Picasso, an autoencoder method that transforms data into an arbitrary predefined 2D shape, e.g., that of an elephant. The authors then compared four kinds of embeddings: the purposefully arbitrary elephant embedding, 2D PCA, *t*-SNE, and UMAP (Figure 1). For this, they used two metrics of embedding quality, both requiring class annotations: *inter-class correlation* measuring how well high-dimensional distances between class centroids are preserved in the 2D embedding and *intra-class correlation* measuring how well class variances are preserved. They found that across three scRNA-seq datasets, 2D PCA performed the best on those metrics, while the elephant embedding scored similar to or better than UMAP and *t*-SNE. We reproduced and confirmed these results (Figure 2a–b).

**Figure 1:**
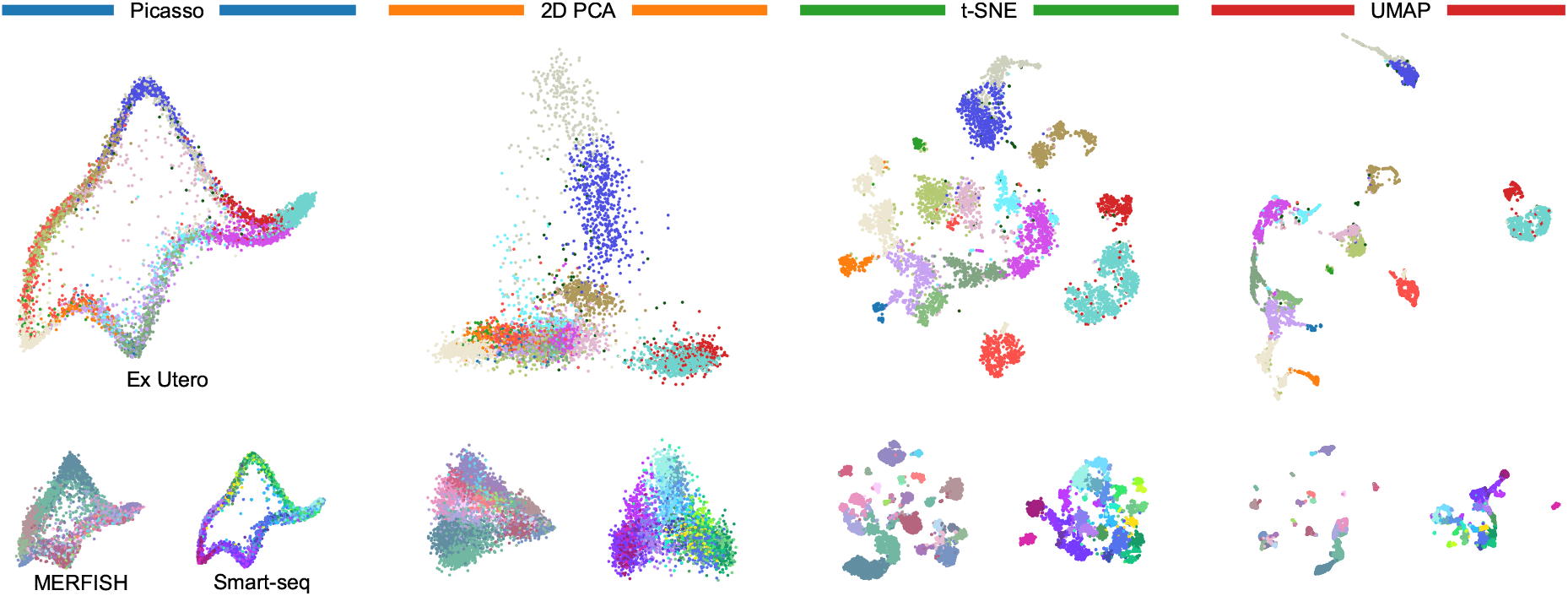
Evaluated embeddings. Large panels: `Ex Utero` dataset. Small panels: `MERFISH` and `Smart-seq` datasets. Colors correspond to cell types and are taken from Chari and Pachter (2023). See Methods for details.

**Figure 2:**
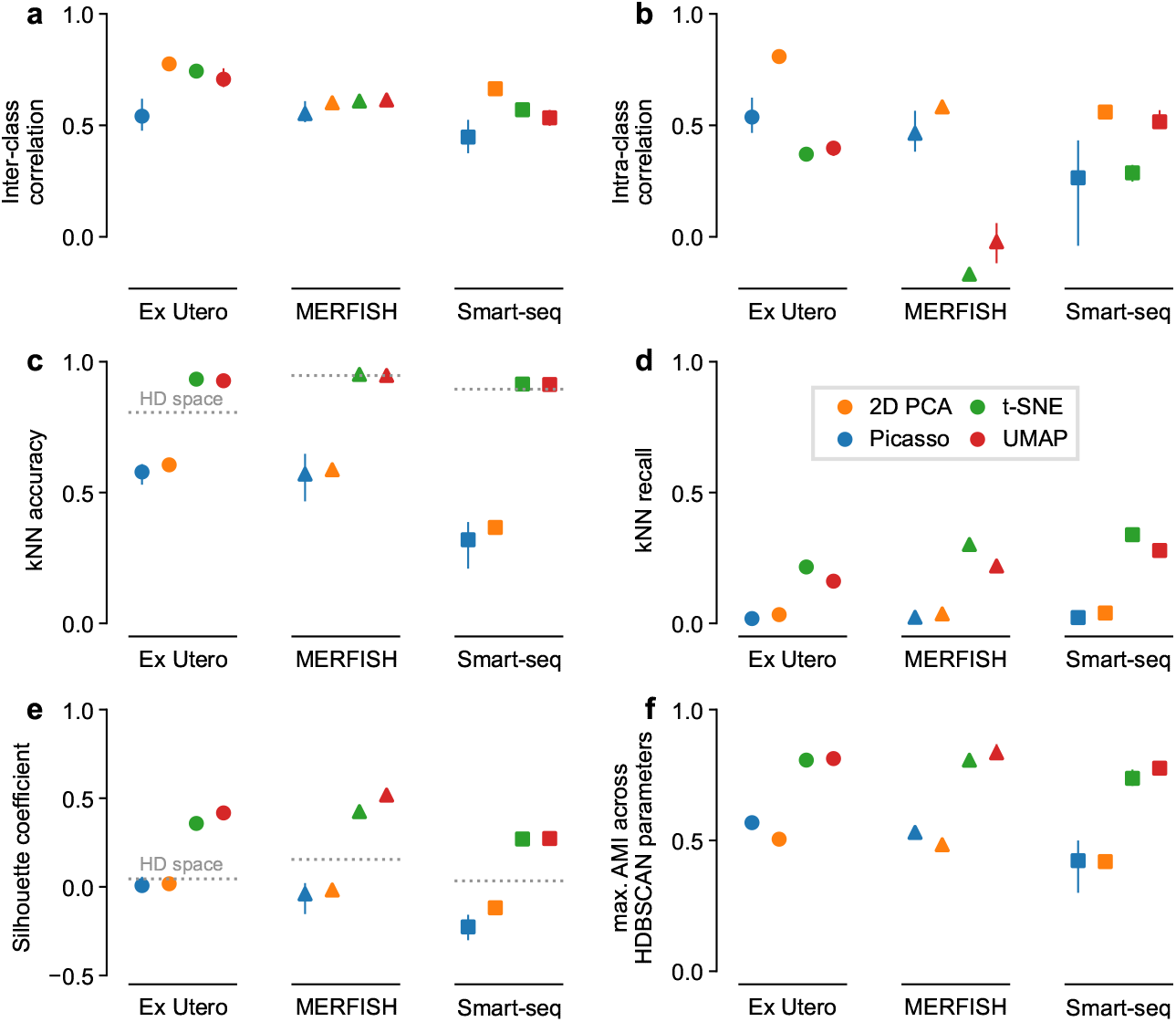
Embedding quality metrics. Panels correspond to metrics, colors correspond to embedding methods, marker shapes correspond to datasets. Averages over five runs, error bars go from the minimum to the maximum across runs. Dotted horizontal lines show the values of the metrics in the high-dimensional gene space. **a–b:** The two metrics from Chari and Pachter (2023), reproducing their Figure 7. **c–d:** *k*NN accuracy and *k*NN recall (*k* = 10). **e:** Silhouette coefficient. **f:** Maximum adjusted mutual information between classes and 2D clusters obtained with HDBSCAN using a range of hyperparameter values.

According to the authors, this means that *t*-SNE and UMAP are as arbitrary and as misleading as the Picasso elephant. Most online discussions and debates about their paper, including posts by the authors themselves, have prominently featured this argument and the powerful elephant metaphor to argue that “it’s time to stop making *t*-SNE & UMAP” plots (Pachter, 2021). In this Comment, we focus exclusively on this argument and do not discuss the rest of the Chari and Pachter (2023) paper.

We believe that this argument is faulty because the metrics used by Chari and Pachter (2023) are insufficient and only quantify a single aspect: both metrics focus on preservation of *distances*, where 2D PCA was unsurprisingly the best. But there is more to embeddings than distance preservation. It is visually apparent in the resulting embeddings that *t*-SNE and UMAP separate cell types, while 2D PCA and Picasso elephant lead to strongly overlapping types (Figure 1), but neither of the two metrics quantified that. Biologists are often interested in cell clusters, and so preservation of cell neighborhoods and visual separation of meaningful cell groups are important properties of 2D embeddings.

To quantify these aspects neglected by Chari and Pachter (2023), we used four additional metrics, commonly employed in benchmark studies (Espadoto et al., 2021; Huang et al., 2022; Wang et al., 2023a): *k*-nearest-neighbor (*k*NN) accuracy, *k*NN recall (Lee and Verleysen, 2009), the silhouette coefficient (Rousseeuw, 1987), and the adjusted mutual information (AMI) between clusters and class labels (Vinh et al., 2009).

The *k*NN accuracy quantifies how often the 2D neighbors are from the same class, while the *k*NN recall quantifies how often the 2D neighbors are the same as the highdimensional neighbors. In both metrics, UMAP and *t*-SNE consistently and strongly outperformed PCA and Picasso elephant embeddings (Figure 2c–d, *>*90% vs. *<*62% accuracy, *>*15% vs. *<*5% recall for all datasets). Even though the *k*NN recall was below 40% for all methods (Figure 2d), *k*NN accuracy was always above 90% for both UMAP and *t*-SNE (Figure 2c). This means that even though UMAP and *t*-SNE are not able to preserve high-dimensional nearest neighbors exactly, the low-dimensional neighbors tend to be from a close vicinity in the high-dimensional space, have the same cell type, and hence allow reliable *k*NN classification. In contrast, 2D PCA and the Picasso elephant fail at that.

The silhouette coefficient and the AMI both evaluate to what extent cell types appear as isolated islands in 2D. Specifically, the silhouette coefficient measures how compact and separated the given classes are in 2D, while the AMI evaluates how well clustering in 2D recovers the classes. In both metrics, *t*-SNE and UMAP strongly outperformed 2D PCA and Picasso elephant embeddings (Figure 2e– f, *>*0.3 difference in silhouette score, *>*0.25 difference in AMI), in agreement with the visual impression (Figure 1). The *k*NN accuracy and the silhouette coefficient can also be computed directly in the high-dimensional gene space.

We found that *t*-SNE and UMAP showed similar or higher *k*NN accuracy and much higher silhouette coefficient than the original high-dimensional space (Figure 2c,e). This suggests that high-dimensional distances suffer from the curse of dimensionality, and that it may in fact be undesirable to preserve them in 2D visualisations. Indeed, single-cell biologists rarely use multidimensional scaling (MDS), an embedding method explicitly designed to preserve distances, because MDS often fails to represent the cluster structure in the data. This further underscores why using only distance-preservation metrics, as Chari and Pachter (2023) did, is misguided.

All presented metrics except *k*NN recall rely on class labels, and our analysis, following Chari and Pachter (2023), used labels derived in original publications via clustering. Therefore, these labels do not necessarily correspond to biological ground truth, and could potentially lead to biased comparisons. To address this concern, we used negative binomial sampling based on the `Ex Utero` dataset to simulate a dataset with known ground truth classes. Analyzing this simulated dataset gave the same conclusions: 2D PCA scored the best in the distance-based correlation metrics of Chari and Pachter (2023), but only *t*-SNE and UMAP could separate the true classes, while Picasso and 2D PCA failed at that (Figure S1).

Taken together, our results point to the elephant in the room: Even though they are not designed to preserve pairwise distances, *t*-SNE and UMAP embeddings are not arbitrary and do preserve meaningful structure of single-cell data, especially local neighborhoods and cluster structure. Claiming that Picasso and *t*-SNE/UMAP are “quantitatively similar in terms of fidelity to the data in ambient dimension” (Chari and Pachter, 2023) is wrong. They are not.

That said, we do agree with Chari and Pachter (2023) that 2D visualisations distort distances and should not be blindly trusted. Moreover, as Chari and Pachter (2023), we do not recommend to use 2D embeddings for any quantitative downstream analysis. However, paraphrasing George Box (Box, 1979), we can say that *all 2D embeddings of high-dimensional data are wrong, but some are useful*. Indeed, one can use 2D embeddings to form hypotheses about the data structure, ranging from data quality control and sanity-checking of any algorithmic output, to more general hypotheses about cluster separability, relationships between adjacent clusters, or presence of outlying clusters. Of course, any generated insight should then be validated in the high-dimensional data by other means. Here, our conclusion differs strongly from that of Chari and Pachter (2023): while they claim that UMAP and *t*SNE are “counter-productive for exploratory […] analyses”, we endorse them for that very purpose.

## Methods

### Datasets and preprocessing

We used the same datasets as Chari and Pachter (2023) (Table 1) and followed the same pre-processing steps. The `Ex Utero` data was already log-normalized. We filtered empty genes and cells, and selected the 2 000 most highly variable genes (HVGs) with `scanpy.pp.highly variable genes()` with default settings (Wolf et al., 2018). The `MERFISH` data was already normalized, and we only performed the `log1p()` transform. The `Smart-seq` data was already log-normalized and had HVGs selected, so we used it as is. Chari and Pachter (2023) additionally performed a standardization step on all datasets, which we omitted for simplicity, as it did not change the result qualitatively (see our Github repository for a direct comparison). Despite small differences in preprocessing choices, we obtained qualitatively very similar results in Figure 2a–b to what the original authors reported in their Figure 7.

**Table 1:** Datasets. `Ex Utero`: Files in GEO accession GSE149372. `MERFISH` and `Smart-seq`: DOIs.

| Name | Cells | Genes | Classes | Source | Count matrix | Metadata |
| --- | --- | --- | --- | --- | --- | --- |
| Ex Utero | 6 205 | 19 588 | 19 | Aguilera-Castrejon et al. (2021) | `normalized.assay85` | `MetaData.85` |
| MERFISH | 6 963 | 254 | 25 | Zhang et al. (2020) | `10.22002/D1.2064` | `10.22002/D1.2063` |
| Smart-seq | 3 850 | 1 999 | 28 | Kim et al. (2019) | `10.22002/D1.2071` | `10.22002/D1.2067` |

### Simulation

We used negative binomial sampling to obtain a simulated version of the `Ex Utero` dataset with known ground-truth classes. For each cluster and each gene *g* in the original dataset, we computed the proportion *p*_*g*_ of UMI counts of this gene among all UMI counts in the cluster. For each cell *c* belonging to this cluster in the original data, we then sampled new counts *X*_*cg*_ ∼ NB(*μ* = *n*_*c*_*p*_*g*_, *θ* = 10), where *n*_*c*_ is the cell’s original total UMI count. Overdispersion parameter *θ* = 10 leads to some additional variance compared to the Poisson distribution. This procedure preserved the number of genes, the number of cells, and all class abundances, and ensured realistic marginal distributions of simulated counts per cell and per gene. The counts of each simulated gene in each class followed an independent negative binomial distribution around the gene’s mean expression in the original `Ex Utero` cluster. Finally, we performed the same pre-processing as above on the simulated counts (depth normalization, scaling normalized counts to 10 000 counts per cell, `log1p()` transform, `scanpy` default HVG selection).

### Embeddings

We used the high-dimensional gene space after pre-processing and gene selection as input to all embedding methods. For the elephant embeddings, we used the original Picasso code by Chari and Pachter (2023) with minimal adjustments needed to provide the random seed for reproducibility (https://github.com/berenslab/picasso). We ran Picasso for 500 epochs with default settings. For PCA, we used `scikit-learn 1.3.0` (Pedregosa et al., 2011) with default parameters. For *t*SNE and UMAP, we followed Chari and Pachter (2023) and first reduced the pre-processed count matrices to 50 dimensions with PCA and used that as input to `openTSNE 1.0.1` (Poličar et al., 2019) and `umap-learn 0.5.5` with default parameters. The 50-dimensional PCA was used in no other part of the analysis. In all plots, we used the class labels and colors from Chari and Pachter (2023), except for minor adjustments to the `Ex Utero` colors, where we introduced four additional colors to make all classes discernible.

### Embedding quality metrics

Following Chari and Pachter (2023), we computed their intra- and inter-class correlation metrics using both *L*^1^ and *L*^2^ distances (see our Github repository for a direct comparison). As we did not observe qualitative differences between the two variants, we only showed *L*^2^ results here, and also used *L*^2^ distances for all other metrics.

For *k*NN accuracy, we used the *k* nearest neighbors in the 2D embedding to predict the class of each cell with a majority vote (this is essentially a leave-one-out crossvalidation procedure). We reported raw accuracy here, but class-balanced accuracy gave qualitatively the same results (see our Github repository). For *k*NN recall, we computed (for each cell) the fraction of the *k* nearest neighbors in the 2D embedding that are also among the *k* nearest neighbors in the high-dimensional space. For both *k*NN metrics, we used *k* = 10, and averaged over all cells.

For the maximum AMI metric, we ran HDBSCAN (McInnes and Healy, 2017) from `scikit-learn` on each embedding for nine hyperparameter values `min samples` = `min size clusters` ∈ *{*5, 10, 15, 20, 30, 40, 50, 75, 100*}*. All points that HDBSCAN left unclustered (noise points) we assigned to their nearest clusters. We then computed the adjusted mutual information (AMI) between each HDBSCAN result and the given cell type class labels, and picked the largest AMI. This way, the best performing hyperparameter was chosen for each embedding and each dataset.

The silhouette coefficient of each cell is defined as (*b* − *w*)*/* max(*b, w*) where *w* is the average distance to cells from the same class and *b* is the average distance to cells in the nearest other class. The silhouette coefficient is then averaged across all cells. We used `scikit-learn` to find *k*NNs, and to compute AMI and silhouette coefficients.

For all metrics that required a high-dimensional reference space for comparison (inter-class and intra-class correlations, *k*NN recall), we used the same high-dimensional gene space that we used as input to the embedding methods.

### Code

Our code in Python is available at https://github.com/berenslab/elephant-in-the-room.

## Acknowledgements

We thank Erik van Nimwegen, Sebastian Damrich, and Pavlin Poličar for discussions. The work was funded by the Deutsche Forschungsgemeinschaft (DFG) (Heisenberg Professorship to PB, BE5601/8-1; Excellence cluster 2064 “Machine Learning — New Perspectives for Science”, EXC 390727645; Excellence cluster 2181 “STRUCTURES”, EXC 390900948), the Gemeinnützige Hertie-Stiftung, the European Union (ERC 101039115 “NextMechMod”), and the National Institutes of Health (UM1MH130981). Views and opinions expressed are however those of the authors only and do not necessarily reflect those of the European Union or the European Research Council Executive Agency. Neither the European Union nor the granting authority can be held responsible for them. The content is solely the responsibility of the authors and does not necessarily represent the official views of the National Institutes of Health.

## Supplementary Figures

**Figure S1:**
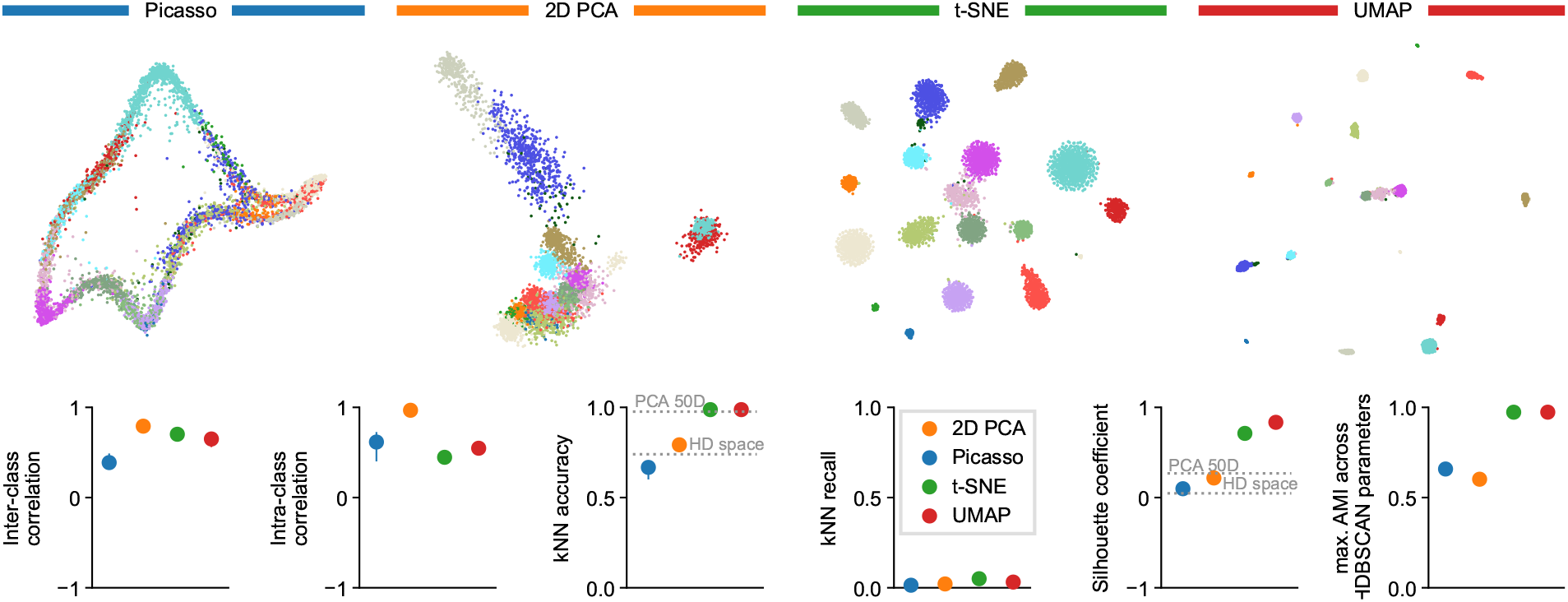
Simulated dataset with ground truth labels. Simulation was based on the `Ex Utero` dataset and generated 19 classes using negative binomial sampling (see Methods for details). Top row: Embeddings as in Figure 1. Bottom row: Embedding quality metrics as in Figure 2. The *k*NN recall values are very low because simulated classes do not have any internal structure. Dotted horizontal lines show the *k*NN accuracy and silhouette score in the high-dimensional gene space (“HD space”) and the 50-dimensional PCA space (“PCA 50D”).

## References

Alejandro Aguilera-Castrejon, Bernardo Oldak, Tom Shani, Nadir Ghanem, Chen Itzkovich, Sharon Slomovich, Shadi Tarazi, Jonathan Bayerl, Valeriya Chugaeva, Muneef Ayyash, et al. Ex utero mouse embryogenesis from pregastrulation to late organogenesis. Nature, 593(7857):119–124, 2021.

Etienne Becht, Leland McInnes, John Healy, Charles-Antoine Dutertre, Immanuel WH Kwok, Lai Guan Ng, Florent Ginhoux, and Evan W Newell. Dimensionality reduction for visualizing single-cell data using UMAP. Nature Biotechnology, 37(1):38–44, 2019.

George EP Box. Robustness in the strategy of scientific model building. In Robustness in statistics, pages 201– 236. Elsevier, 1979.

Tara Chari and Lior Pachter. The specious art of single-cell genomics. PLOS Computational Biology, 19(8):e1011288, 2023.

Mateus Espadoto, Rafael M Martins, Andreas Kerren, Nina ST Hirata, and Alexandru C Telea. Toward a quantitative survey of dimension reduction techniques. IEEE Transactions on Visualization and Computer Graphics, 27(3):2153–2173, 2021.

Haiyang Huang, Yingfan Wang, Cynthia Rudin, and Edward P Browne. Towards a comprehensive evaluation of dimension reduction methods for transcriptomic data visualization. Communications Biology, 5(1):719, 2022.

Dong-Wook Kim, Zizhen Yao, Lucas T Graybuck, Tae Kyung Kim, Thuc Nghi Nguyen, Kimberly A Smith, Olivia Fong, Lynn Yi, Noushin Koulena, Nico Pierson, et al. Multimodal analysis of cell types in a hypothalamic node controlling social behavior. Cell, 179(3):713–728, 2019.

Dmitry Kobak and Philipp Berens. The art of using t-SNE for single-cell transcriptomics. Nature Communications, 10(1):5416, 2019.

John A Lee and Michel Verleysen. Quality assessment of dimensionality reduction: Rank-based criteria. Neurocomputing, 72(7-9):1431–1443, 2009.

Leland McInnes and John Healy. Accelerated hierarchical density based clustering. In 2017 IEEE International Conference on Data Mining Workshops (ICDMW), pages 33–42. IEEE, 2017.

Leland McInnes, John Healy, and James Melville. UMAP: Uniform manifold approximation and projection for dimension reduction. 1802.03426, 2018.

Luis Gustavo Nonato and Michael Aupetit. Multidimensional projection for visual analytics: Linking techniques with distortions, tasks, and layout enrichment. IEEE Transactions on Visualization and Computer Graphics, 25(8):2650–2673, 2018.

Lior Pachter, 2021. URL https://web.archive.org/web/20240729115631/ https://archive.is/2024.07.29-115414 https://x.com/lpachter/status/1431325969411821572.

Fabian Pedregosa, Gaël Varoquaux, Alexandre Gramfort, Vincent Michel, Bertrand Thirion, Olivier Grisel, Mathieu Blondel, Peter Prettenhofer, Ron Weiss, Vincent Dubourg, et al. Scikit-learn: Machine learning in Python. The Journal of Machine Learning Research, 12:2825– 2830, 2011.

Pavlin G Poličar, Martin Străzar, and Blăz Zupan. openTSNE: a modular python library for t-SNE dimensionality reduction and embedding. BioRxiv, page 731877, 2019.

Peter J Rousseeuw. Silhouettes: a graphical aid to the interpretation and validation of cluster analysis. Journal of Computational and Applied Mathematics, 20:53–65, 1987.

Laurens Van der Maaten and Geoffrey Hinton. Visualizing data using t-SNE. Journal of Machine Learning Research, 9(11), 2008.

Nguyen Xuan Vinh, Julien Epps, and James Bailey. Information theoretic measures for clusterings comparison: is a correction for chance necessary? In Proceedings of the 26th Annual International Conference on Machine Learning, pages 1073–1080, 2009.

Kaiwen Wang, Yuqiu Yang, Fangjiang Wu, Bing Song, Xinlei Wang, and Tao Wang. Comparative analysis of dimension reduction methods for cytometry by time-offlight data. Nature Communications, 14(1):1836, 2023a.

Shu Wang, Eduardo D Sontag, and Douglas A Lauffenburger. What cannot be seen correctly in 2D visualizations of single-cell ‘omics data? Cell Systems, 14(9):723–731, 2023b.

F Alexander Wolf, Philipp Angerer, and Fabian J Theis. SCANPY: large-scale single-cell gene expression data analysis. Genome Biology, 19:1–5, 2018.

Meng Zhang, Stephen W Eichhorn, Brian Zingg, Zizhen Yao, Hongkui Zeng, Hongwei Dong, and Xiaowei Zhuang. Molecular, spatial and projection diversity of neurons in primary motor cortex revealed by in situ single-cell transcriptomics. BioRxiv, pages 2020–06, 2020.

